# Constrained dynamics of oligonucleotides in the phase-separated droplets

**DOI:** 10.1101/2023.02.04.527127

**Authors:** Anupam Singh, Shashi Thutupalli, Manoj Kumar, Sandeep Ameta

## Abstract

Phase-separated droplets are excellent means of compartmentalizing functional molecules and have been shown as excellent models for protocells. Although complex functions based on oligonucleotides have been studied, we still lack an understanding of how the oligonucleotide dynamics are affected by the condensed internal environment of these droplets. Particularly, we lack high-resolution experimental measurements of the dynamical parameters that control oligonucleotide diffusion inside the phase-separated droplets. In addition, there is no clarity on how these dynamical parameters differ in the charged (coacervates) *vs* non-charged (aqueous two-phase system, ATPS) environment of these droplets. In this study, using fluorescence correlation spectroscopy (FCS), we demonstrate the constrained dynamics of oligonucleotides inside membraneless phase-separated droplets at an unprecedented resolution. We further compare transport properties at different lengths of oligonucleotides as well as salt concentrations. We observe that among all the parameters the oligonucleotide’s caging (spatial restriction in the movement) inside the matrix has a considerable impact on the diffusive dynamics. Our study provides a way of unravelling, quantifying and understanding physical parameters governing the polymer transport dynamics inside the phase-separated droplets.

## Introduction

Compartmentalization of biomolecules is an indispensable step in constructing synthetic protocells from the bottom-up approaches and is crucial to several origin-of-life scenarios (1–3). Among different ways of compartmentalization, protocells based on liquid-liquid phase separation are of particular interest as they have been shown to support different oligonucleotide-based catalytic functions (4–10) and their permeable nature enables implementation of out-of-equilibrium protocols by allowing for dynamical exchange with the environment (11–14). Despite the demonstration of numerous oligonucleotide-based catalytic functions, it is unclear what dynamical parameters govern oligonucleotide diffusion in the condensed environment of these phase-separated droplets. For example, the caging phenomenon, in which molecules are spatially trapped in a polymer mesh and exhibit non-Brownian behavior, restricts the movement of oligonucleotides within the compartment (15). It has been shown that such cages or micro-environment affects the gene expression (16), consequently, in the context of protocells, the constrained movement of oligonucleotide due to caging may affect its catalytic function or interaction with other components. Therefore, it is crucial to experimentally measure the constrain dynamics, such as caging, cross-over time, and overall diffusivity of the oligonucleotides within the heterogeneous internal environment of protocells. Such dynamics can be assessed using microscopy-based methods such as FRAP (fluorescence recovery after photobleaching) and FCS (fluorescence correlation spectroscopy), where the former quantifies lateral diffusion of molecules following spatial bleaching of the sample and, the latter relies on time-correlation of diffusing biomolecules to construct its diffusion trajectory. Although FRAP is frequently employed to assess the system’s diffusive character (17) and has also been used to understand DNA dynamics inside coacervates (18), it lacks the resolution for analyzing the dynamic behavior of molecules at faster timescales. In contrast, FCS measurements display a variety of timescales, with the faster ones (10 – 100 *μ s*) corresponding to the polymer’s dynamics in its cage and the slower ones to its translational diffusion in the compartment (19). Additionally, FCS can also resolve dynamics at smaller length scales (~ 10 *nm*) corresponding to the cage dimensions. This enables the extraction of crossover time (*i.e*., the amount of time it takes for a molecule to move from one cage to another) and dynamics within the cage, which can be utilized to obtain an approximation of caging effect (19–23).

In this study, using FCS, we measure various diffusion-related parameters influencing the oligonucleotide dynamics inside the condensed environment of two different types of phase-separated droplets, non-charged ATPS (aqueous two-phase based system, based on segregative interactions) and charged coacervates (based on associative interaction between polymers). These two compartment systems differ in terms of internal packing, cage dimensions, charge density, and interaction with encapsulated oligonucleotides, which may affect catalysis and diffusion dynamics. We also compare these quantities for different DNA lengths and salt concentrations. The results show that due to differences in the caging effect, DNA oligonucleotides are more diffusive inside the ATPS droplets than charged coacervates, demonstrating that caging constraints play a significant role in controlling the diffusive dynamics.

## Results and Discussion

We used polyethylene glycol-Dextran (PEG-DEX) ATPS and poly(diallyldimethylammonium chloride)-adenosine triphosphate (PDAC-ATP) coacervate droplets as a representatives of non-charged and charged phase-separated compartments, respectively (Fig. 1a). To study the polymer dynamics using FCS, a fluorescently labeled 20nt DNA oligonucleotide was also co-encapsulated in these phase separated droplets (5’-Alexa Flour 488, Material and Methods, Supplementary Table I). As expected, PDAC-ATP droplets were much efficient in partitioning the oligonucleotide inside compared to PEG-DEX droplets (Supplementary Fig. 1). Then timedependent fluorescence intensity fluctuations of the labeled oligonucleotides inside each phase-separated compartment (number of droplets, *n* ~ 25 per condition) were used to obtain correlation curves. These correlation curves were used to calculate the physical parameters such as diffusivity (*D*), minimum of the local exponent of the MSD (*α_min_*), cage cross-over time (*τ_α_*), and mean squared displacement within the cage (*MSD_α_*) using a previously established strategy (19, 24–26) (Fig. 1a, b, Material and Methods).

**Fig. 1.**
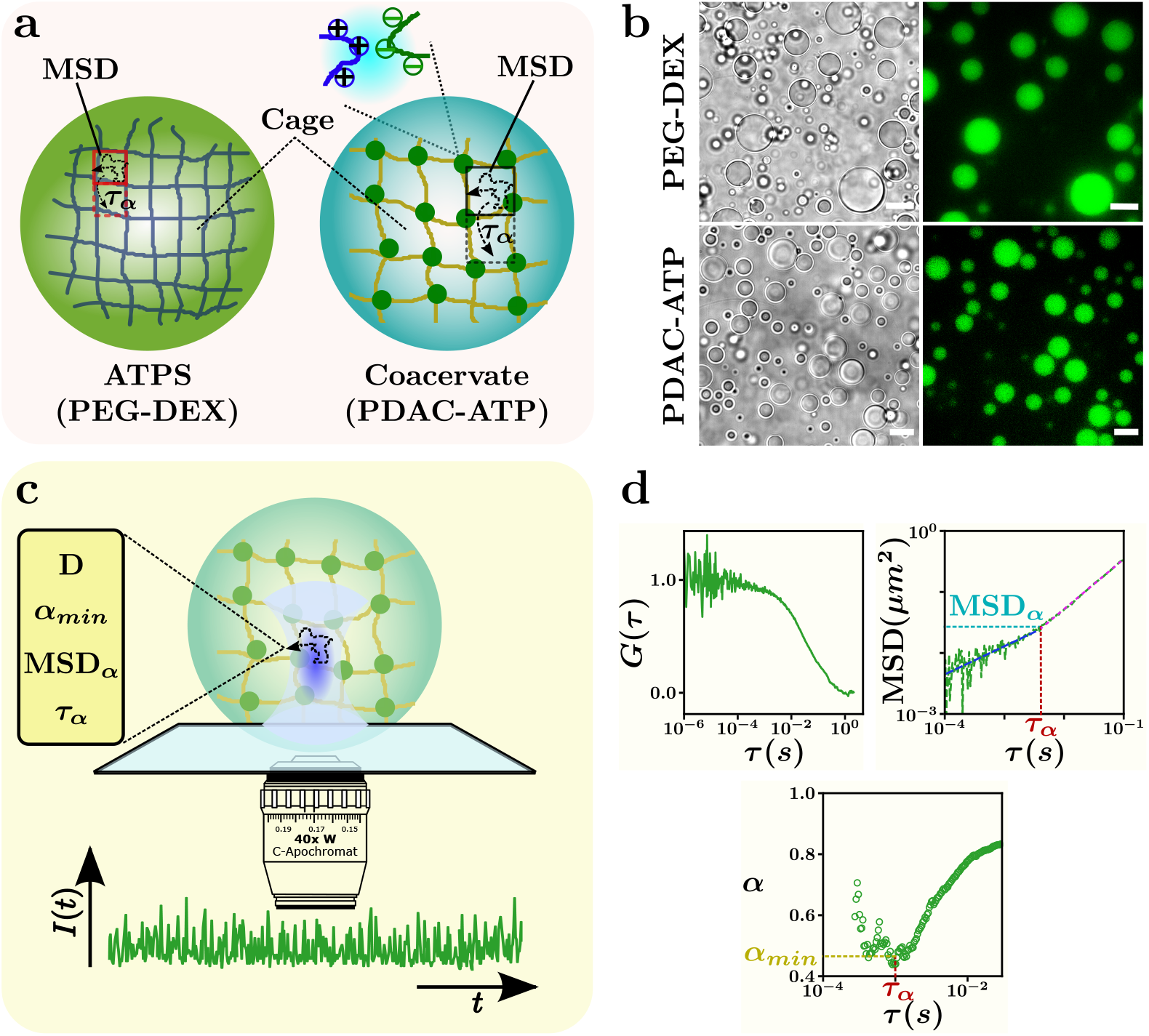
Droplet system and FCS measurement strategy. **(a)** Schematic showing the phase-separated droplets, the hypothetical internal structure (cage), and hoping of oligonucleotide between the cages. **(b)** Microscopy images of PEG-DEX (*top*) and PDAC-ATP (*bottom*) droplets visualized in bright-field channel (*left*) and fluorescence channel (*right*). Fluorescence is due to Alexa Flour 488 labeled 20nt DNA oligonucleotide co-encapsulated in the droplets. PEG-DEX system forms larger droplets compared to the PDAC-ATP ones. The scale bar is 10 *μm* for PDAC-ATP and 20 *μm* for PEG-DEX droplets. **(c)** Schematic showing the strategy for obtaining oligonucleotide dynamics and parameters D, *α_min_*, *MSD_α_* and *τ_α_*. To derive correlation plots, fluorescence intensity (*I*(*t*)) overtime profiles are measured. **(d)** The normalized auto-correlation (*G*(*τ*)) curves obtained from FCS measurements (top *left*) which are utilized to obtain the MSD *vs τ* plots (*top right*). These plots are then further analyzed to extract the local exponent plot to obtain cross-over dynamics of oligonucleotides along with the caging (*bottom*).

The parameter *D* represents the observed molecular diffusion on slower time scales, whereas parameters *α_min_*, *τ_α_*, and *MSD_α_* describe the dynamics of oligonucleotides at faster time scales and are critical in observing constrained dynamics of biomolecules due to internal structuration of the phase-separated droplets (Fig1c, d). The local exponent ‘*α*’ computed from *MSD* represents a sub-diffusive or diffusive behaviour of oligonucleotide in the droplet. While the molecules in phase-separated droplets appear diffusive (*α* ~ 1) on slower time scales, the sub-diffusive regime (*α* < 1) can be identified at faster time scales representing the crowding in the surrounding environment (Fig1d). The change in *α* (*α* < 1 → *α* = 1) can be used to describe the transition from a sub-diffusive regime to a diffusive regime, where the smaller values of “*a_min_*” indicates the effect of increased local crowding. Such sub-diffusive regions in our system may form as a result of interactions between the polymeric components in the charged system and due to polymer demixing in non-charged systems. As a result, the DNA oligonucleotides would be caged, limiting their movement within the phase-separated droplet. The amount of time needed for the molecules to relax from their caged condition is determined by the cross-over time (*τ_α_*) corresponding to the *α_min_*. Similarly, *MSD_α_* (= *MSD*(*τ_α_*)) represents the mean squared displacement within the cage before transitions to a regime where the molecules diffuse freely and can be treated as a proxy for the cage’s apparent free area for the oligonucleotide. Therefore, the cage dimensions of two droplet systems can be compared by observing *τ_α_* and *MSD_α_* simultaneously.

At first, using FCS, we compare the dynamics of a 20nt long DNA oligonucleotide in both of the droplet systems. The results show that the oligonucleotide is more diffusive in the ATPS system compared to coacervates (4.58 *vs* 0.95 *μm*^2^/*s*, Fig. 2a, and Supplementary Table II). While oligonucleotide behave sub-diffusively at faster timescales in both systems (*α_min_* < 1), the values of *α_min_* does not vary significantly between the two (Fig. 2b). The higher diffusivity in PEG-DEX droplets suggests a less constrained system, which could be caused by small cages through which oligonucleotides can move faster, or by stronger interactions between oligonucleotides and polymer in PDAC-ATP droplets. Interestingly, the cross-over time (*τ_α_*) for ATPS droplets is shorter than for coacervates (2.81 × 10^−4^ *s vs* 2.69 × 10^−3^*s*, Fig. 2c, Supplementary Table II), which, together with the lower *MSD_α_* values (Fig. 2d), strongly suggests a smaller cage size with weaker caging effect in the ATPS droplets.

**Fig. 2.**
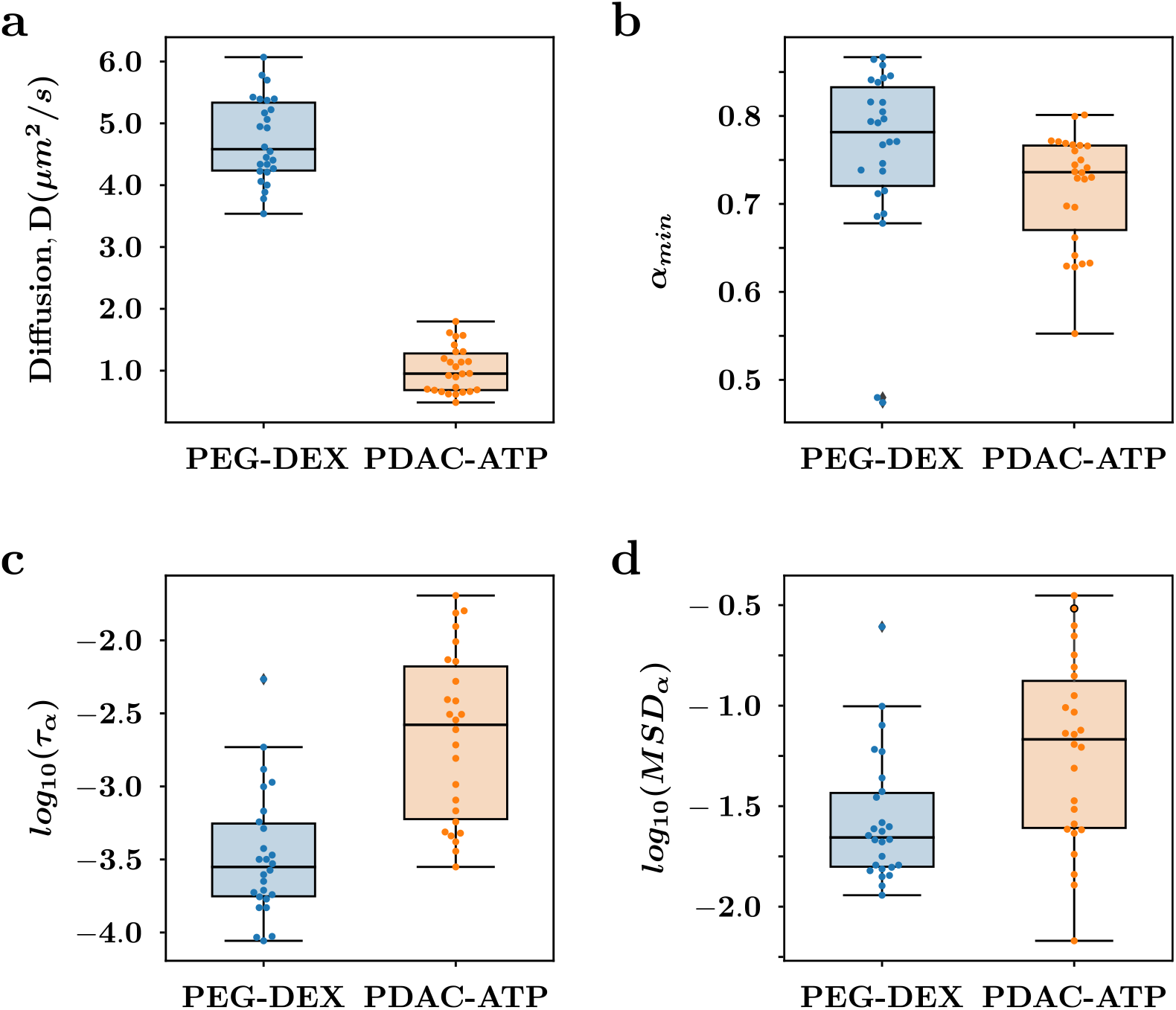
Comparison of physical parameters for a 20nt DNA oligonucleotide. **(a)** Graph showing the comparison of diffusivity (*D*) of a 20nt oligonucleotide in the PEG-DEX (*blue*) and PDAC-ATP (*orange*) droplets. **(b)** Measurements of minima of the local exponent *α* (*α_min_*) representing the crowdedness of the surrounding (inside the phase-separated droplets) in the sub-diffusive regime. **(c)** Comparison of the logarithm of cross-over timescale obtained from *α_min_* suggesting the time required for the molecule to relax from the cage. **(d)** Comparison of logarithm of mean squared displacement (*MSD_α_*, calculated using the *α_min_*) of 20nt DNA oligonucleotide within the cage.

Such weaker caging effect might be caused by either loosely packed polymers, or reduced interactions between the probe (oligonucleotide) and caging polymers (18). These findings imply that the dynamics of the oligonucleotide within these phase-separated droplets are largely governed by the internal mesoscopic structural parameters, such as caging, allowing DNA oligonucleotide to diffuse more quickly in ATPS droplets compared to coacervates.

Another factor that may have an impact on the diffusion dynamics is the charge density of the encapsulated molecules, particularly in the charged environment of coacervate droplets. The charge density can be easily altered by varying the length of the oligonucleotides. In order to determine all the parameters (*D*, *α_min_*, *τ_α_*, and *MSD_α_*), we then measured the dynamics of 40nt and 80nt DNA ins-die these droplets and compare them with parameters vales obtained for 20nt DNA (Fig. 3). We found that, for both droplet systems, the diffusivity reduced significantly as the length of oligonucleotide increases. This change is higher in PEG-DEX (~ 74% reduction; from *D* = 4.58*μm*^2^/*s* for 20nt to *D* = 1.18*μm*^2^/*s* for 80 nt) than in PDAC-ATP coacervates (~ 48%reduction; from *D* = 0.9 *μm*^2^/*s* to *D* = 0.49*μm*^2^/*s*, Fig. 3a, Supplementary Table III & IV). Similarly, there was a significant change in the *α_min_*, which was found to be more affected in PEG-DEX droplets than in coacervate droplets. However, unlike for 20nt DNA, the *α_min_* for 80nt DNA was lower in the PEG-DEX system compared to coacervates (0.335 vs 0.498, Fig. 3b, Supplementary Table III & IV), despite the PEG-DEX system having slightly higher diffusivity *D* for 80nt. This indicates that the bulkier oligonucleotides are more constrained and have lower diffusivity. The other two parameters, *τ_α_* and *MSD_α_*, were found to be less affected in the PEG-DEX system, however in PDAC-ATP droplets, the *MSD_α_* was more affected (Fig. 3c, d). This demonstrates that even if the cage cross-over time in the coacervates is faster, there will be a lesser free space inside the cage for the 80nt DNA than there is in PEG-DEX droplets. These findings further indicate that the oligonucleotide dynamics in these phase-separated droplets are primarily influenced by the caging.

**Fig. 3.**
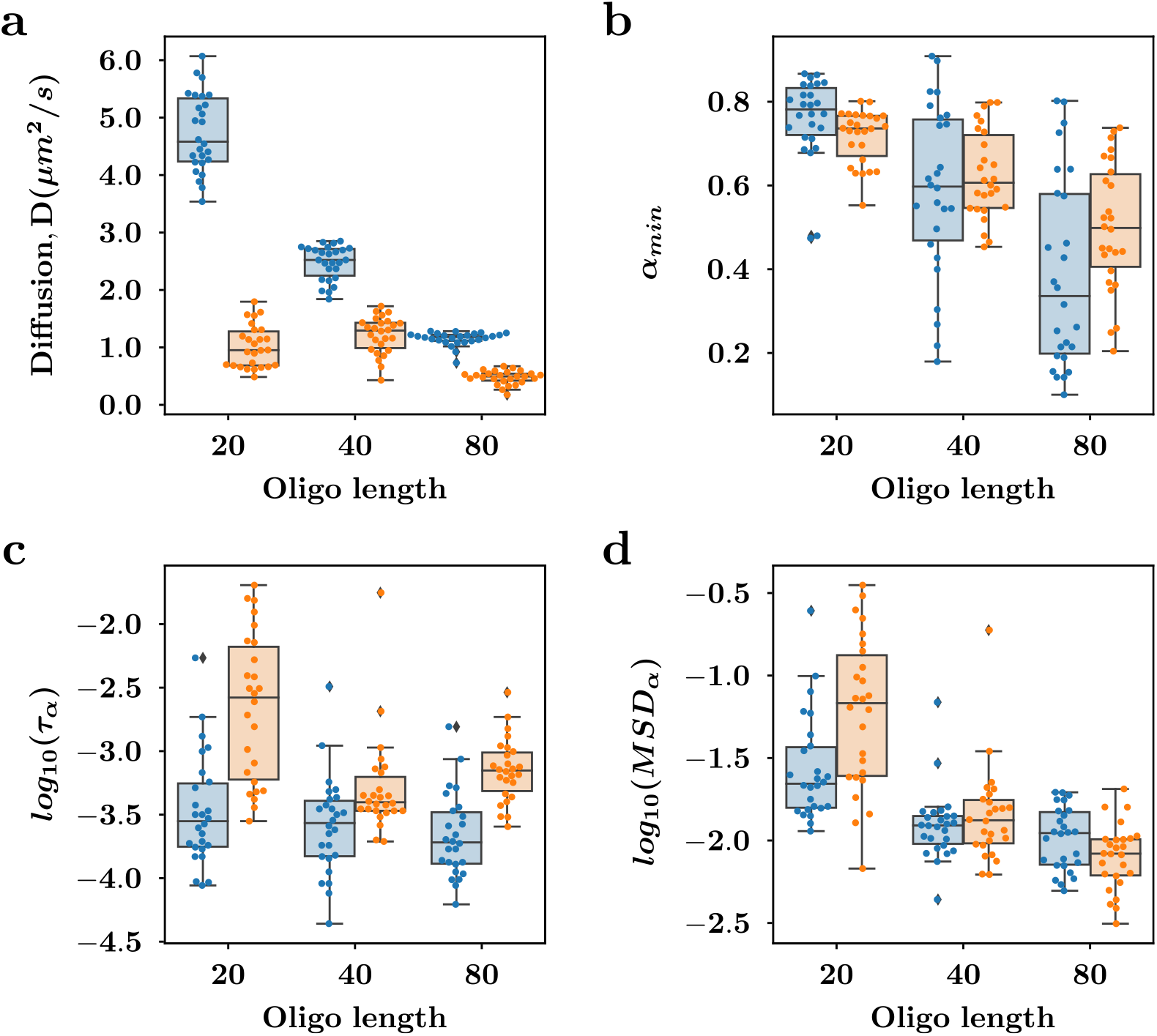
FCS measurements using different length of oligonucleotides. **(a)** Graph showing the diffusivity comparison of different length oligonucleotides (20nt, 40nt and 80nt) in PEG-DEX (*blue*) and PDAC-ATP (*orange*) droplets. **(b)** Measurements of minima of the local exponent *α* (*α_min_*) for different lengths oligonucleotides to determine the effect of cage size on the oligonucleotide dynamics. **(c)** Comparison of the logarithm of cross-over timescale obtained from *α_min_* showing the time required for the oligonucleotides of different lengths to relax from the cage of PDAC-ATP and PEG-DEX system. **(d)** Comparison of logarithm of mean squared displacement (*MSD_α_*) of 20nt, 40nt and 80nt oligonucleotide within the cage of PEG-DEX and PDAC-ATP system.

The internal dynamics and stability of protocells can also be affected by the constant external environmental pressure. The external environment can be altered using salt which can shield negatively charged oligonucleotides as well as the charged polymers (in the case of PDAC-ATP droplets). To test this, we prepared droplets of PDAC-ATP and PEG-DEX in the presence of various monovalent salt concentrations (0 – 50 *mM* NaCl). Although, we also prepared the droplets in presence of divalent salt (MgCl_2_), PDAC-ATP droplets were found to be stable only below a salt concentration of 2.5 *mM*, therefore not proceeded further. Our results show that the salt has a stronger effect on PDAC-ATP droplets than on PEG-DEX system. The turbidity, number density, and the size of PDAC-ATP droplets have significantly reduced, whereas PEG-DEX droplets were largely remains unaffected in the presence of salt (Fig. 4a, b). We then analyzed these droplets to probe the effect of salt on *D*, *α_min_*, *τ_α_*, and *MSD_α_* (Fig. 4c-f). For both droplet systems, the parameters *α_min_*, *τ_α_*, and *MSD_α_* did not change significantly with salt concentrations (Fig. 4d-f). However, there was a significant increase in *D* for PEG-DEX droplets, whereas no change in *D* was observed for PDAC-ATP droplets despite the notable effect on size (Fig. 4b, c). This indicates that for PDAC-ATP droplets, salt doesn’t change the internal environment of droplets but rather accelerates the dissolution process. The formation of coacervates and the internal environment are a function of polymer length, charge density, and as well as the concentration of the polymers. As in the current study we do not change the polymer length, concentration or charges, the electroneutrality point should not change. Furthermore, as reported earlier, associative phase-separated droplets dissolve when the salt is added above a critical concentration that depends on type of the salt as well as density inside the droplet phase (27). In our case, it is most likely that after above 25 *mM* NaCl, PDAC-ATP droplets are dissolving without affecting the electroneutrality point (28) or internal structuration (i.e. cage dimensions). Contrarily, for PEG-DEX droplets, the increase in *D* with an increase in salt concentration is surprising as there was no effect on the droplet size, number density and other dynamical parameters (Fig. 4b-f). The only charged component in PEG-DEX system is oligonucleotide itself, therefore the charge shielding by salt could selectively affect diffusivity. In addition, PEG is also known to act as dehydrating agent (29), which may change the water content inside the droplets affecting the mesoscopic properties.

**Fig. 4.**
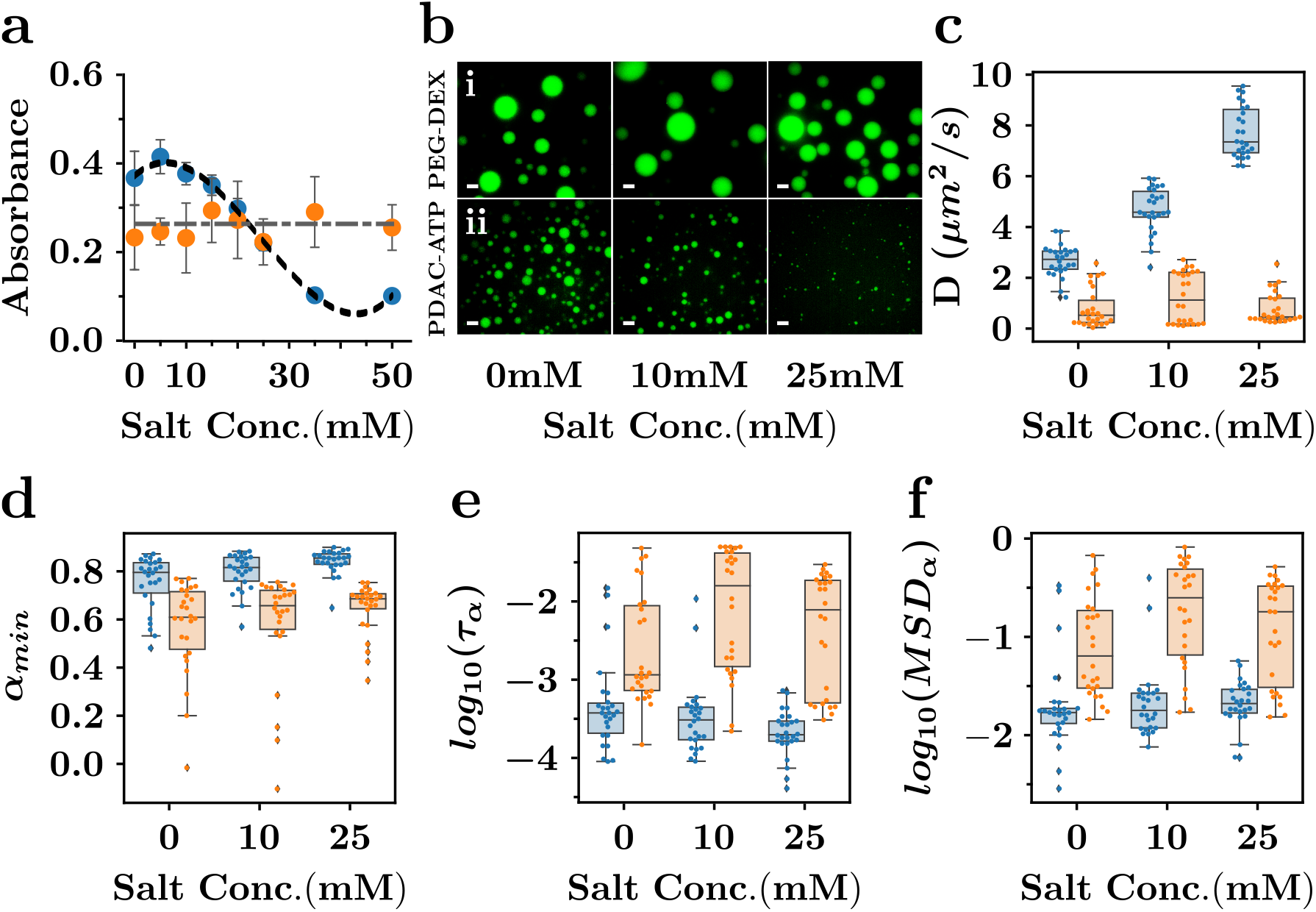
Effect of salt on the dynamic of DNA oligonucleotide inside phase-separated droplets. **(a)** Graph showing the turbidity measurements of samples with PEG-DEX and PDAC-ATP droplets prepared in presence of different NaCl concentrations. Turbidity was quantified by measuring the absorbance of the solution at 420 *nm* wavelength. All the measurements were performed in triplicates and mean absorbance with standard deviation is plotted here. **(b)** Fluorescence microscopy image of PEG-DEX and PDAC-ATP droplets in presence of different NaCl concentrations. Fluorescence is due to the encapsulation of labeled 20nt DNA (Alexa Flour 488, same as used above). The scale bars for PDAC-ATP and PEG-DEX are 10 *μM* and 20 *μM*, respectively, **(c)** Graph showing the effect of salt on the diffusivity of 20nt oligonucleotide DNA in PEG-DEX (blue) and PDAC-ATP (orange) system. **(d)** Measurements of *α* (*α_min_*) at different salt concentrations on the PEG-DEX (blue) and PDAC-ATP (orange) system. **(e)** Logarithm of cross-over timescale obtained from *αmin* at different salt concentrations to determine the time required for the oligonucleotides to relax from the cage of PDAC-ATP and PEG-DEX system. **(f)** The comparison of the logarithm of mean squared displacement (*MSD_α_*) of 20nt oligonucleotide within the cage of PEG-DEX and PDAC-ATP system at different salt concentrations.

## Conclusion

The development of an evolvable protocell capable of undergoing cycles of growth and division is of paramount importance (11). Understanding the dynamics of information-containing polymers inside such protocells is very crucial, especially when the internal environment is heterogeneous, *e.g*. in phase-separated droplets. The proposition of such droplets as model “protocell” by Oparin and Haldane (30, 31) has sparked a number of experimental works involving functional DNA/RNA and phase-separated droplets (5–9). However, there are limited studies examining the internal dynamics of oligonucleotides inside these droplets. In particular, we lack systematic and high-resolution experimental data on the dynamical behavior of nucleic acids inside the condensed environment of these droplets. There have been attempts to understand RNA or DNA dynamics using FRAP or FCS (17, 18), however, in those studies the oligonucleotide was also used as a component that contributes to phase separation, and thus its diffusion is coupled to the compartment itself. In this regard, the current study, to the best of our knowledge, is one of the first ones to use FCS to explore the dynamics of an oligonucleotide that is enclosed inside a phase-separated droplet, enabling measurements of dynamics with a very high spatiotemporal resolution. By changing the charge density of DNA (DNA length) as well as the surrounding environment (salt), we systematically examined the dynamics of two distinct phase-separated systems (ATPS and coacervates). The findings show that the diffusive dynamics is significantly influenced by the interior structure of the droplet, such as caging. In both systems, the oligonucleotide movement was sub-diffusive at faster timescales (*α* < 1) indicating that the environment is crowded and constrained. Despite this, smaller cages in PEG-DEX droplets (the lower values of *τ_α_* and *MSD_α_*) allow oligonucleotide to move faster (higher *D*). In the case of PDAC-ATP, even an increase in charge density did not decrease the diffusivity (*D*) significantly, suggesting that charge interaction is not the major contributor in affecting the movement. However, it would be interesting to measure the dynamics of even longer oligonucleotides (higher charge density) in order to check if there is a trade-off between the effect of caging and the oligonucleotide’s charge density. Furthermore, measuring the differences in the dynamics caused due to structured oligonucleotides versus non-structured oligonucleotides (used in this study) will also be helpful in creating a comprehensive understanding of the dynamics of nucleic acids inside the phase-separated compartments.

## Materials and Methods

### A. Materials

All the experiments were performed using DNAse/RNase free water (Thermo Fisher Scientific Product No. 10977015). Chemicals were purchased from Sigma-Aldrich unless specified otherwise: PDAC (Poly(diallyldimethylammonium chloride), Sigma-Aldrich, Product no. 409022), ATP (Adenosine 5’-triphosphate disodium salt, Sigma-Aldrich, Product no. A7699), PEG (polyethylene glycol, Mw~ 8000, Sigma-Aldrich, Product no. P2139), Dextran(Mw~ 9000-11000, Sigma-Aldrich, Product no. D9260. DNA oligonucleotides were obtained from Integrated DNA Technologies (IDT) and are mentioned in Supplementary Table I.

### B. Methods

#### B.1. Preparation of phase-separated droplets

For all experiments, phase-separated droplets were prepared in 50 —100 *μL* scale volume by mixing the polymers along with DNA oligonucleotide. **PEG-DEX droplets**: For preparing PEG-DEX ATPS, we mix 2.5 *μL* of respective DNA oligonucleotide (100 *nM* stock) with 10 *μL* of dextran (30(*w/v*)%) polymer which then mixed with 10 *μL* of PEG (50(*w/v*)%) polymer. The PEG-DEX solution was thoroughly mixed using a vortex-mixer to ensure homogeneous phase separation throughout the sample. When prepared with salt, respective amount of salt (NaCl) was added to the dextran solution prior to addition of PEG polymer solution. **PDAC-ATP droplets**: Coacervate samples were prepared in 50 *μL* volume with 1 *nM* DNA oligonucleotide (100 *nM* stock), 50 *mM* Tris-Cl buffer pH 8.0, 10 *mM* ATP in water and were mixed thoroughly using pipette. Then 10 *mM* of PDAC was added to the solution and mixed using vortex resulting in turbid solution of coacervates. When prepared with salt, respective amount of NaCl was added prior to addition of PDAC. To facilitate FCS measurement and imagining, all the DNA oligonucleotides were ordered with 5’-modification of Alexa Flour 488 fluorophore (from IDT DNA, Supplementary Table I).

#### B.2. FCS on coacervates

The single point FCS measurements were performed as described earlier by Singh *et al*. and a brief description of the experimental setup, equipment, calibration and analysis are further discussed below (24–26, 32).

##### Experimental Setup and equipment

FCS experiments were performed on the Confocal setup (Carl Zeiss LSM 780) operated in photon counting mode, objective: C-Apochromat 40x 1.2 NA W Corr M27; excitation sources: 488 *nm* (Ar-ion laser: ~ 1 *μW*); emission range for detectors: 499 – 552 *nm*; pinhole diameter: 37 *μm;* temperature: 23.0 ± 1.0°C]. The time-intensity traces and the autocorrelations were recorded and obtained from Zen blue 10 software from Carl Zeiss.

##### Dynamical physical quantities calculated from FCS

DNA based oligonucleotides labeled at 5’-end were used to measure their constraint dynamics phase-separated droplets using FCS. The time-intensity traces of the fluctuating end of oligonucleotides were obtained, and auto-correlated to extract correlation *G*(*τ*) at lag-time *τ*. The fluctuations of oligonucleotide end correspond to the diffusion of the molecules in the caged environment which can be studied using mean squared deviation (〈*r*^2^(*τ*)〉) of the end of the polymer by applying appropriate mathematical transformation and solving for the roots of the expression:

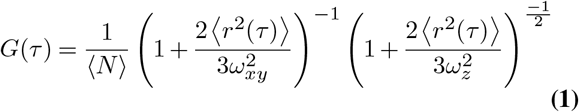

where 〈*N*〉, *ω_xy_* and *ω_z_* are the number of molecules in the confocal volume, the lateral and the axial radius of the confocal volume respectively. 〈*r*^2^(*τ*〉) is obtained by solving the Eqn.1 and obtaining the roots by Broyden’s good method (33,34) with the help of a custom written python code on normalized *G*(*τ*) and using the pre-determined values of *ω_xy_* and *ω_z_*. *G*(0) is the mean of first 10 lag-time values of *G*(*τ*) with which *G*(*τ*) is normalized, resulting in 〈*N*〉 = 1.

The dynamics of the oligonucleotides in caged and freely diffusing regime were obtained from 〈*r*^2^(*τ*)〉 by separating the two regimes by identifying the local slope *α*(*τ*).

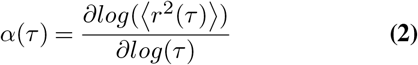

*α*(*τ*) was numerically calculated by taking data points from 〈*r*^2^(*τ*)〉 corresponding to 1 order of magnitude of lag-time (*i.e*. 66 data-points in current setup). The minima of *α*(*τ*) [*α_min_* = min (*α*(*τ*))] corresponds to the cross-over from caged dynamics to a freely diffusing oligonucleotide. The *α_min_* was calculated using a custom written code which finds the minimum and takes the average of nearest 5 *α*(*τ*), and then the 〈*α*(*τ_α_*)〉 is further plotted and analysed. Further, the cross-over time *τ_α_* is the lag-time corresponding to *α_min_*, the mean squared displacement of the cross-over from caged dynamics to free diffusion *MSD_α_* = 〈*r*^2^(*τ_α_*)〉.

##### Determining confocal volume for FCS

The confocal beam expansions *ω_xy_* and *ω_z_* in equatorial and axial planes were calculated using Alexa Flour 488 with known diffusion *D* = (414 *μm*^2^/*sec*) at 25°C (35). 10 FCS curves were recorded for Alexa Flour 488 alone in solution using the same way as described. Then using the expression 1, mean diffusion time *τ_D_* = 34.75 ± 3.7 *μs* and structure parameter *ρ* = 6.49 were calculated. Further, *ω_xy_*(~ 0.240 *μm*) is obtained by using known diffusion *D_A488_* and calculated *τ_D_* in expression:

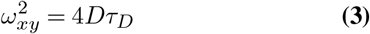

and the axial expansion *ω_z_* = *ω_xy_* × *ρ*.

##### Experiments to determine dynamical physical parameters

The experiments were carried out in custom made PDMS (polydimethylsiloxane) wells bonded on the corrected glass coverslip (175 ± 5 *μm*). The PEG-DEX droplets did not wet the surface, however the charged PDAC-ATP droplets perfectly wet the glass surface. Therefore, to avoid droplet wetting on glass surface, the coverslips were coated with a thin layer (20 *μl*) of optical glue (Norland Optical Adhesive 81) by spin coating at 2000 *rpm* for 30 *s* and exposing it under UV for 30 *s*. For non-charged ATPS droplets, glass coverslips were directly used without any modification. After adding droplets, PDMS wells were sealed with coverslip and subjected to FCS measurements as described above. For each droplet, ~ 12 FCS curves with 30 *s* each were recorded and in total 26 droplets were measured for each condition. The aberrant curves—which differed by more than one standard deviation and are caused by oligonucleotide aggregation—were manually eliminated during the post-processing. Furthermore, in order to remove any anomaly in data coming from drifting of droplets, the fluorescence intensity of FCS were to eliminate the data of very slow timescales, and by visually comparing droplet images before and after FCS measurements.

## Acknowledgement

We thank Jyoti Prasad Banerjee (NCBS Bangalore) for the feedback on the manuscript. We thank the Central Imaging and Flow Cytometry Facility (CIFF) at the NCBS. Author also thanks laboratory of Dr. Marino Zerial (MPI-CBG, Dresden) for providing support.

## Supplementary Information

The supplementary information contains tables and additional results.

