## Supplementary Information for "Constrained dynamics of oligonucleotides in the phase-separated droplets"

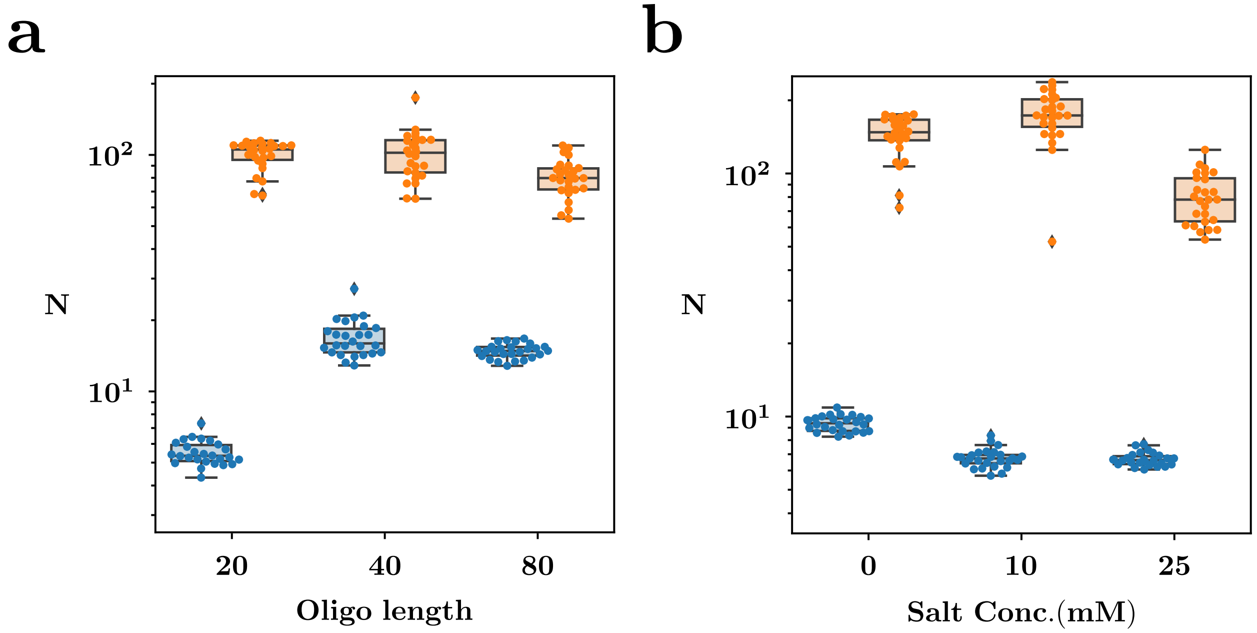


FIG. S1. **Number of DNA oligonucleotide molecules encapsulated.** Graph showing the number of DNA molecules in the confocal volume of the PEG-DEX (*blue*) and PDAC-ATP (*orange*) droplets. Number of molecules are shown here as a function of oligonucleotide length (**a**) and salt concentration (**b**). The boxplot shows the mean, first and second quartile from each experiment. The sample size for each case is $\approx 25$

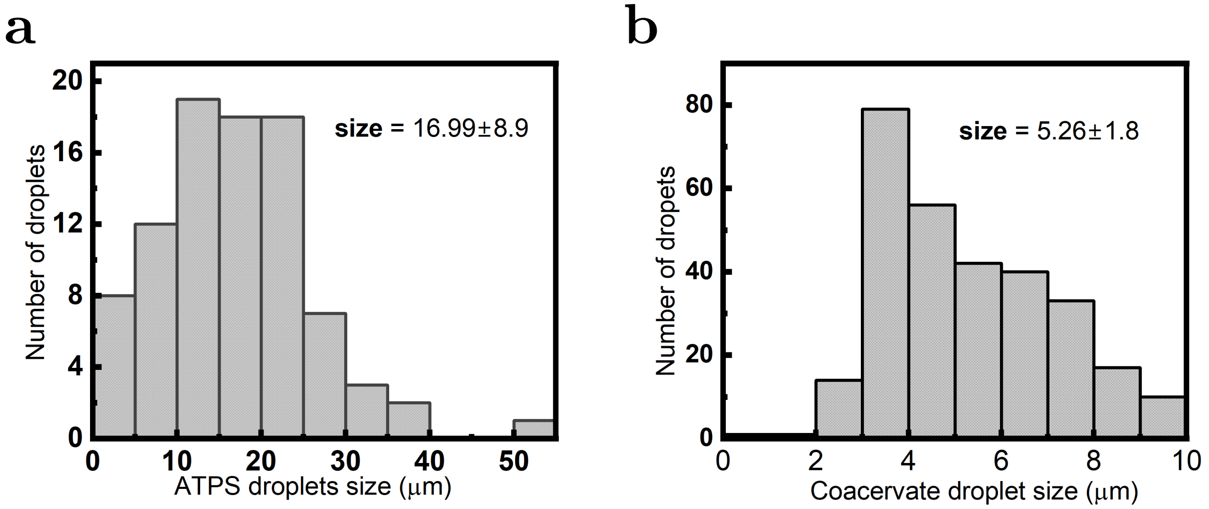


FIG. S2. **Droplet size distribution.** The size distribution of the PEG-DEX (**a**) and PDAC-ATP droplets (**b**). The average size for PEG-DEX and PDAC-ATP droplets is $16.99\mu m$ (sample size = 88) and $5.26\mu m$ (sample size = 293), respectively. The sizes were calculated using the '*analyse particle*' function in Fiji software [1].

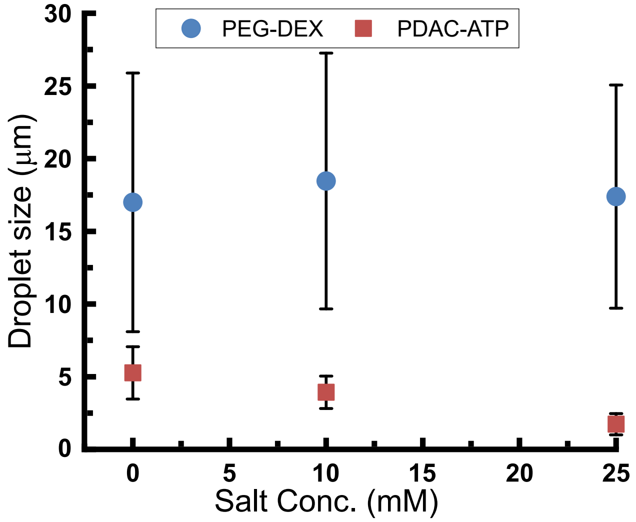


FIG. S3. **Effects of salt concentration on droplet size.** The droplet size distribution for PEG-DEX (blue) and PDAC-ATP droplets (red) under different salt concentrations (0 $mM$, 10 $mM$ and 25 $mM$ NaCl). The average size for PEG-DEX droplet remains similar while varying the salt concentration. However, the PDAC-ATP droplets size reduced with an increase in salt concentration. The droplet size was calculated using the '*analyse particle*' function in Fiji software [1]. The sample size for 0 $mM$, 10 $mM$ and 25 $mM$ NaCl in PEG-DEX system is 88, 17 and 53 respectively, and for PDAC-ATP coacervates is 293, 300 and 90 respectively.

TABLE I. **Fluorescently labelled oligonucleotides sequences and their length used in the study.**

| **Oligo sequence** | **Oligo length** |
|---|---|
| TGT TTT GTG TTG GGT GGG TT/3AlexF488N/ | 20 |
| GTG TGT GTG TGT GGT TTG TGT GTT TTG TGT TGG GTG GGT T/3AlexF488N/ | 40 |
| TG TGT GTG TGT GGG GTG TGT GTG TGT GTG GGT TTT TTT GTG TGT GTG TGT GTG GTT TGT GTG TTT TGT GTT GGG TGG GTT/3AlexF488N/ | 80 |

TABLE II. **Comparison of dynamical properties of 20mer oligonucleotides in charged *vs* non-charged system (sample size = 25 *per condition*).**

| Quantity | PDA-ATP_20mer | PEG-DEX_20mer |
|---|---|---|
| **D** ($\mu m^2/s$) | 0.95 | 4.58 |
| $\alpha_{min}$ | 0.73 | 0.78 |
| $\tau_c$ ($\times 10^{-3} s$) | 2.69 | 0.281 |
| **MSD** ($\times 10^{-2} \mu m^2$) | $6.76 \times 10^{-2}$ | 2.23 |

TABLE III. **The effect of oligonucleotide length on localization and dynamical properties in the PDAC-ATP droplets (sample size = 25 *per condition*).**†

| Quantity | 20mer | 40mer | 80mer |
|---|---|---|---|
| $\langle \mathbf{N} \rangle$ | $105.411^{+109.636}_{-95.298}$ | $102.143^{+115.449}_{-84.288}$ | $79.819^{+87.701}_{-71.435}$ |
| $\mathbf{D}$ $(\mu m^2/s)$ | $0.951^{+1.278}_{-0.685}$ | $1.293^{+1.428}_{-0.987}$ | $0.494^{+0.543}_{-0.421}$ |
| **MSD** $(\times 10^{-2}\ \mu m^2)$ | $6.80^{+13.30}_{-2.23}$ | $1.32^{+1.76}_{-0.96}$ | $0.833^{+1.018}_{-0.615}$ |
| $\tau_c$ $(\times 10^{-3} s)$ | $2.642^{+6.637}_{-0.598}$ | $0.396^{+0.628}_{-0.339}$ | $0.704^{+0.977}_{-0.485}$ |
| $\alpha$ | $0.736^{+0.766}_{-0.670}$ | $0.606^{+0.720}_{-0.546}$ | $0.498^{+0.627}_{-0.405}$ |

† Notation $\mathbf{X}^{\mathbf{a}}_{\mathbf{b}}$: $\mathbf{X}$ is the median, $^{\mathbf{a}}$ is the upper quartile and $_{\mathbf{b}}$ is the lower quartile of the distribution.

TABLE IV. **The effect of oligonucleotide length on localization and dynamical properties in the PEG-DEX droplets (sample size = 25 *per condition*).**[†]

| **Quantity** | **20mer** | **40mer** | **80mer** |
|---|---|---|---|
| $\langle \mathbf{N} \rangle$ | $5.341^{+5.934}_{-5.074}$ | $15.958^{+18.398}_{-14.622}$ | $14.825^{+15.416}_{-14.171}$ |
| **D** $(\mu m^2/s)$ | $4.582^{+5.335}_{-4.236}$ | $2.522^{+2.713}_{-2.249}$ | $1.181^{+1.219}_{-1.114}$ |
| **MSD** $(\times 10^{-2}\ \mu m^2)$ | $2.20^{+3.68}_{-1.57}$ | $1.23^{+1.40}_{-0.95}$ | $1.11^{+1.49}_{-0.71}$ |
| $\tau_c\ (\times 10^{-3} s)$ | $0.281^{+0.557}_{-0.177}$ | $0.271^{+0.407}_{-0.148}$ | $0.191^{+0.331}_{-0.130}$ |
| $\alpha$ | $0.781^{+0.832}_{-0.720}$ | $0.597^{+0.757}_{-0.468}$ | $0.335^{+0.579}_{-0.198}$ |

[†] Notation $\mathbf{X}^{\mathbf{a}}_{\mathbf{b}}$: $\mathbf{X}$ is the median, $^{\mathbf{a}}$ is the upper quartile and $_{\mathbf{b}}$ is the lower quartile of the distribution.

TABLE V. **The effect of salt concentration (NaCl) on localization and dynamical properties of 20mer oligonucleotides in PDAC-ATP droplets (sample size=25 *per condition*).†**

| **Quantity** | **0mM NaCl** | **10mM NaCl** | **25mM NaCl** |
|---|---|---|---|
| $\langle \mathbf{N} \rangle$ | $147.788^{+166.418}_{-136.954}$ | $173.252^{+201.905}_{-155.564}$ | $78.079^{+95.571}_{-63.533}$ |
| **D** $(\mu m^2/s)$ | $0.526^{+1.112}_{-0.236}$ | $1.123^{+2.214}_{-0.175}$ | $0.457^{+1.197}_{-0.380}$ |
| **MSD** $(\times 10^{-2}\ \mu m^2)$ | $6.39^{+18.5}_{-3.01}$ | $24.77^{+48.977}_{-6.54}$ | $18.03^{+32.88}_{-3.06}$ |
| $\tau_c\ (\times 10^{-3} s)$ | $1.153^{+8.830}_{-0.729}$ | $15.885^{+41.591}_{-1.472}$ | $7.816^{+18.620}_{-0.505}$ |
| $\alpha$ | $0.609^{+0.715}_{-0.476}$ | $0.657^{+0.721}_{-0.558}$ | $0.685^{+0.706}_{-0.644}$ |

† Notation $\mathbf{X}^{\mathbf{a}}_{\mathbf{b}}$: $\mathbf{X}$ is the median, $^{\mathbf{a}}$ is the upper quartile and $_{\mathbf{b}}$ is the lower quartile of the distribution.

TABLE VI. **The effect of salt concentration (NaCl) on localization and dynamical properties of 20mer oligonucleotides in PEG-DEX droplets (sample size = 25 *per condition*).†**

| **Quantity** | **0mM NaCl** | **10mM NaCl** | **25mM NaCl** |
|---|---|---|---|
| $\langle \mathbf{N} \rangle$ | $9.385^{+9.837}_{-8.730}$ | $6.724^{+6.954}_{-6.421}$ | $6.614^{+6.871}_{-6.373}$ |
| **D** ($\mu m^2/s$) | $2.725^{+3.049}_{-2.337}$ | $4.581^{+5.401}_{-4.389}$ | $7.349^{+8.629}_{-6.924}$ |
| **MSD** ($\times 10^{-2}\ \mu m^2$) | $1.70^{+1.87}_{-1.31}$ | $1.78^{+2.68}_{-1.18}$ | $2.10^{+2.93}_{-1.67}$ |
| $\tau_c$ ($\times 10^{-3} s$) | $0.376^{+0.502}_{-0.207}$ | $0.306^{+0.445}_{-0.171}$ | $0.198^{+0.295}_{-0.164}$ |
| $\alpha$ | $0.795^{+0.835}_{-0.709}$ | $0.815^{+0.857}_{-0.759}$ | $0.855^{+0.872}_{-0.829}$ |

† Notation $\mathbf{X}^{\mathbf{a}}_{\mathbf{b}}$: $\mathbf{X}$ is the median, $^{\mathbf{a}}$ is the upper quartile and $_{\mathbf{b}}$ is the lower quartile of the distribution.

### REFERENCES

---


*
†
‡



[1] J. Schindelin, I. Arganda-Carreras, E. Frise, V. Kaynig, M. Longair, T. Pietzsch, S. Preibisch, C. Rueden, S. Saalfeld, B. Schmid, *et al.*, Nature methods **9**, 676 (2012).
